# Structural insights into mechanisms of zinc scavenging by the *Candida albicans* zincophore Pra1

**DOI:** 10.1101/2025.01.09.632233

**Authors:** Johanna L Syrjanen, Alexandre Nore, Elena Roselletti, Tanmoy Chakraborty, Rajika L Perera, Duncan Wilson

## Abstract

*Candida albicans* causes more than 400,000 life-threatening, and half a billion mucosal infections annually. In response to infection, the host limits availability of essential micronutrients, including zinc, to restrict growth of the invading pathogen. As assimilation of zinc is essential for *C. albicans* pathogenicity, its limitation induces the secretion of the zincophore protein Pra1 to scavenge zinc from the host. Pra1 also plays a number of important roles in host-pathogen interactions and is conserved in most fungi. However, the structure of fungal zincophores is not known. Here, we present the first cryogenic electron microscopy structures of *C. albicans* Pra1 in its apo- and zinc-bound states, at 2.8 and 2.5 Å resolution respectively. Our work reveals a hexameric ring-like assembly with multiple zinc binding sites. Through genetic studies, we show that one of these zinc binding sites is essential for *C. albicans* growth under zinc restriction but does not affect the inflammatory properties of Pra1. These data provide a foundation for future work to explore the structural basis of Pra1-mediated host-pathogen interactions, *C. albicans* zinc uptake, as well therapeutics development.

## Introduction

*Candida albicans* is one of the most common fungi in humans, typically living commensally within the oral cavity, gastrointestinal and urogenital tracts^1^. However, overgrowth of *C. albicans* can cause infections, which in otherwise healthy individuals commonly present as oral or vaginal thrush (candidiasis) and in the immunocompromised can lead to life-threatening invasive candidiasis of internal organs or bloodstream infection (candidemia)^2^. The ability of *C. albicans* to capture zinc from its environment is critical to its pathogenicity^3^. During infection, the host employs a strategy called nutritional immunity to restrict the availability of essential trace elements to the invading pathogen^4^. However, under conditions of low zinc availability, *C. albicans* secretes the zinc scavenging (’zincophore’) protein Pra1 to capture zinc from the host^3^. As well as capturing zinc for the fungus, we have recently shown that Pra1 is responsible for the inflammatory immunopathology of vulvovaginal candidiasis and has multiple immune-modulatory functions^5,6^. Although the physiochemical properties of individual Pra1 peptide fragments have been investigated, the mechanisms of Pra1 zinc sequestration remain unexplored on a structural level ^7,8^. In this study, we present the first cryo-EM structures of *C. albicans* Pra1 in the apo-state as well as in the zinc-bound state at 2.8 Å and 2.5 Å resolution, respectively. Our study reveals a hexameric, wheel-like structure, with key zinc binding sites on the exterior rim of the wheel. Further, we show that genetic mutations in *C. albicans* carrying mutations of these sites show severely impeded growth under low zinc conditions but does not affect Pra1 immuno-stimulatory properties. Together, our structural and genetic discoveries suggest a molecular mechanism for zinc capture by Pra1. Our work is foundational for further structural, biochemical and genetic studies unravelling the molecular steps in the zincophore activity of Pra1. Because of the central role that Pra1 plays in *C. albicans* pathogenicity, it is a key target for the development of new therapeutics. These structures are a substantial step forward in aiding these efforts.

## Results

We determined the structure of recombinantly expressed *C. albicans* Pra1 to 2.5 Å resolution (Extended Figure 1). The structure reveals a hexameric ring-like assembly (Figure 1a) in which each subunit is related to its neighbour by a 180° rotation (Figure 1b). This leads to two distinct inter-subunit interfaces (Figure 1b-f). From examination of these interfaces, we suggest that the hexameric assembly formed by Pra1 subunits comprises a trimer of dimers. To form a dimer, the N-terminus of a subunit hugs across the main body of its neighbour like an outstretched arm or brace (Figure 1b-d). As illustrated in Figure 1d, these interactions are primarily mediated by salt bridges (Arg34 with Asp71; Asp36 with Arg75; Arg75 with Asp107) and hydrogen bonding (Tyr31 with Arg75; Trp33 with Asp230; Trp37 with the backbone oxygen of Ala106) of the N-terminal brace with its neighbouring subunit. In addition to the extensive N-terminal interactions with the neighbouring subunit, the polar interaction of Ser93 with Glu188 contributes to the intra-dimer interactions (Figure 1e). In contrast to the intra-dimer interactions, the inter-dimer interactions are almost solely driven by polar interactions that involve residues Thr50, Glu54, Thr57, Tyr152, Glu155, Thr154, and Ser159 (Figure 1f, inset).

**Figure 1.**
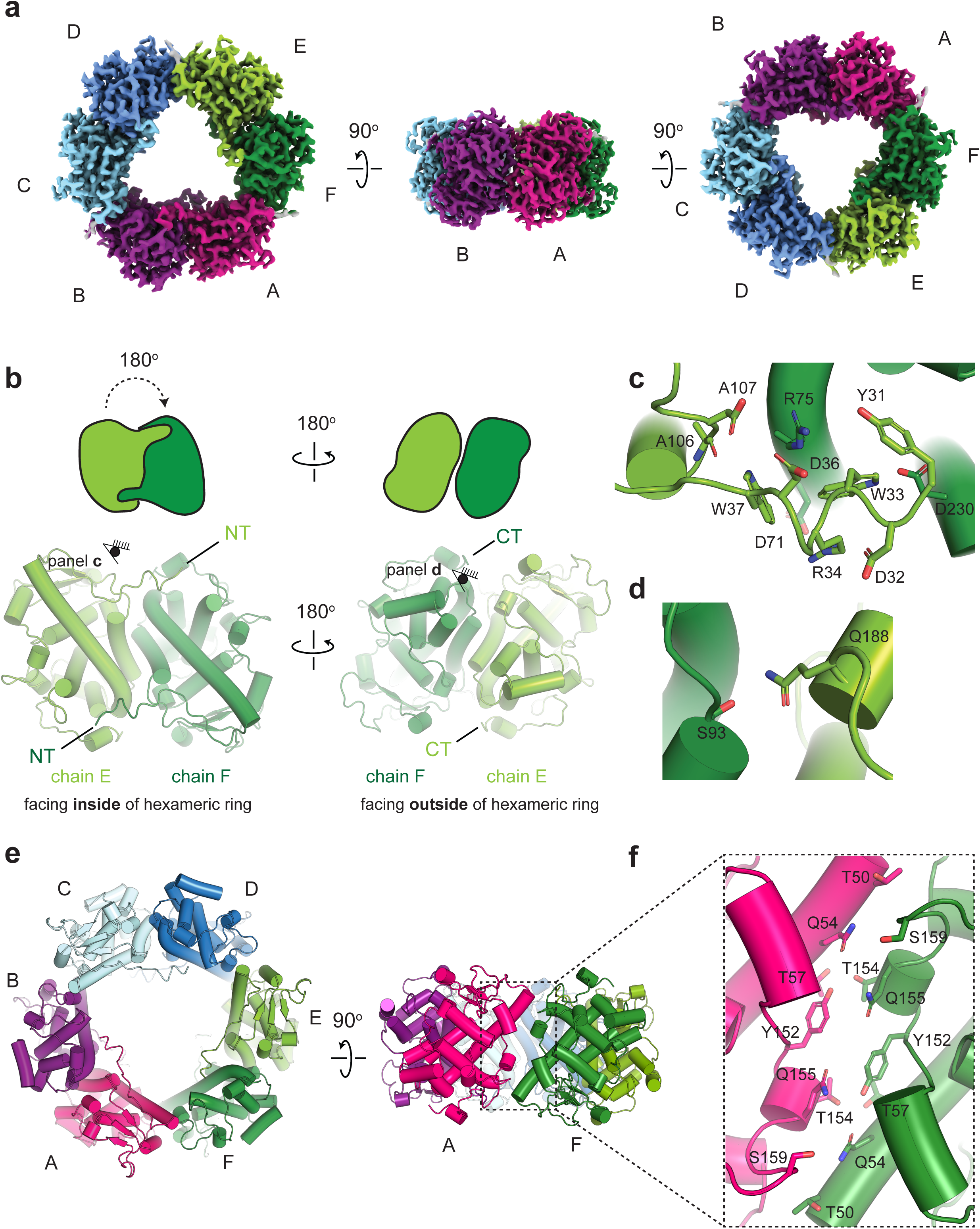
The cryo-EM structure of Pra1 reveals a hexameric assembly. a) The cryo-EM map of *C. albicans* Pra1. b) The hexamer consists of an assembly of a trimer of dimers. A dimer is shown in smudge green (chain E) and forest (chain F). Each Pra1 subunit is related to its neighbour by a 180° rotation, resulting in two distinct interaction interfaces per subunit. Chains E+F, A+B, and C+D make up the dimers. c) The N-terminus of a Pra1 subunit forms a brace or arm that is held across part of its neighbouring subunit to form a dimer. The N-terminal brace intra-dimer interactions are primarily mediated by salt bridges and hydrogen bonds. d) Apart from the N-terminal brace interactions, Q188 and S93 form polar interactions across the intra-dimer interface. e) The interactions at the interdimer interface are polar in nature.

Pra1 is 299 amino acids in length (including its N-terminal signal peptide) and our cryo-EM structure allows the visualisation of residues 31-251, as well as some post-translational modifications. In agreement with original observations of *C. albicans* fungal culture, Pra1 is extensively glycosylated^9^. Asn48, Asn89, Asn135 and Asn208 all showing evidence of N-linked glycosylation. The glycosylated residues are all located in close proximity to subunit interfaces (Figure 2a, Extended Figure 2). Asn48 is close to the intra-dimer interface (Extended Figure 2). In contrast, Asn89, Asn208 and Asn135 are all located close to the inter-dimer interface, leading to a total of six glycosylation sites in this area (Extended Figure 2).

**Figure 2.**
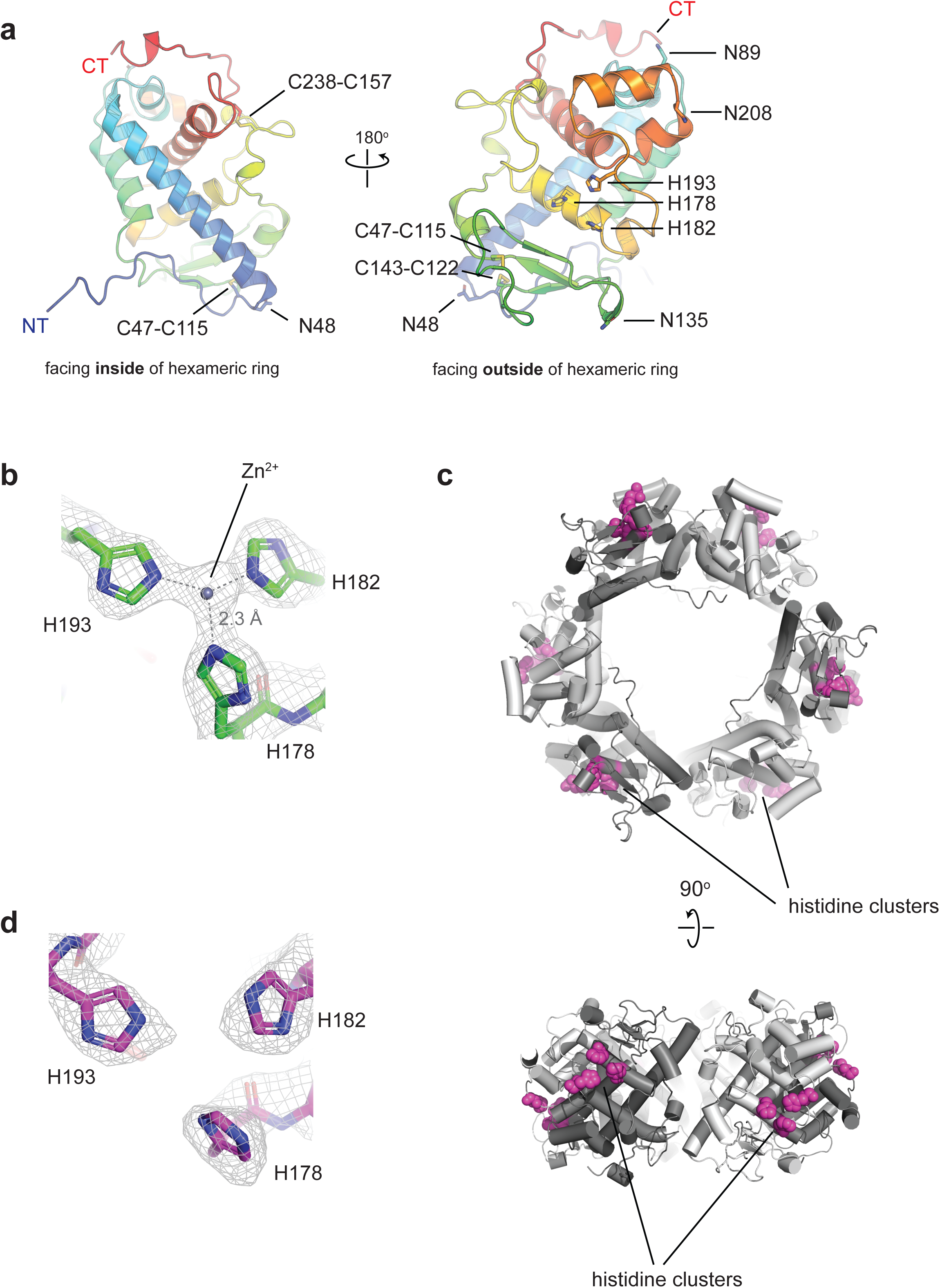
A triad of histidine residues forms a zinc binding site on a Pra1 subunit on the exterior of the hexameric ring. a) The N-terminal and C-terminal ends of the protein are on opposite sides of Pra1. A single subunit consists of several short helices, as well as a long helix 24 amino acids in length and a small beta-sheet. In addition, a Pra1 subunit harbours three disulphide bridges, four N-linked glycosylation. b) A close-up of the histidine triad comprised of His 178, His 182 and His 193 in the presence of 1 mM ZnSO_4_ reveals cryo-EM map density consistent with a metal coordinated by the triad. In this subunit (chain B), the distance from the centre of the zinc to the centre of the τ nitrogen in the imidazole ring of the histidine residue is 2.3 Å for all three histidine side chains. c) The locations of the histidine triad consisting of His178, His 182 and His 193 on the exterior ring are indicated in magenta space fill. Each subunit harbours one histidine triad, leading to six zinc binding sites on the exterior of the Pra1 ring. d) A close-up of the histidine triad in MES pH 6.0 reveals a lack of metal binding under these conditions, as indicated by the absence of continuous cryo-EM density between the three histidines, in contrast to panel b.

Examination of a single subunit shows that the N- and C-termini are located on opposite faces of the protomer (Figure 2a). The 180° rotational relationship between neighbouring subunits means that each of the “top” and the “bottom” faces of the hexameric ring has three N-termini and C- termini. The core of the protomer is made up of five short helices, one longer helix and one beta-sheet (Figure 2a). The longest helix (residues 52-76) spans 24 amino acids across the length of the subunit and faces the interior of the hexameric ring (Figure 2a). The beta-sheet, consisting of three beta-strands (within residues 112-142), is on the exterior of the hexameric ring (Figure 2a). Each Pra1 subunit also has three intramolecular disulphide bonds, all in close proximity to the inter-dimer interface (Figure 2a). Indeed, the overall fold of a Pra1 protomer is similar to that of HEXXH + D type metalloproteases (Extended Figure 3a; a structural superposition of Pra1 with deuterolysin from *Aspergillus oryzae;* PDB code 1EB6, yields an RMSD =3.74 Å). However, the catalytically active glutamic acid residue characteristic of metalloproteases has been replaced with an arginine in Pra1 (Extended Figure 3a).

**Figure 3.**
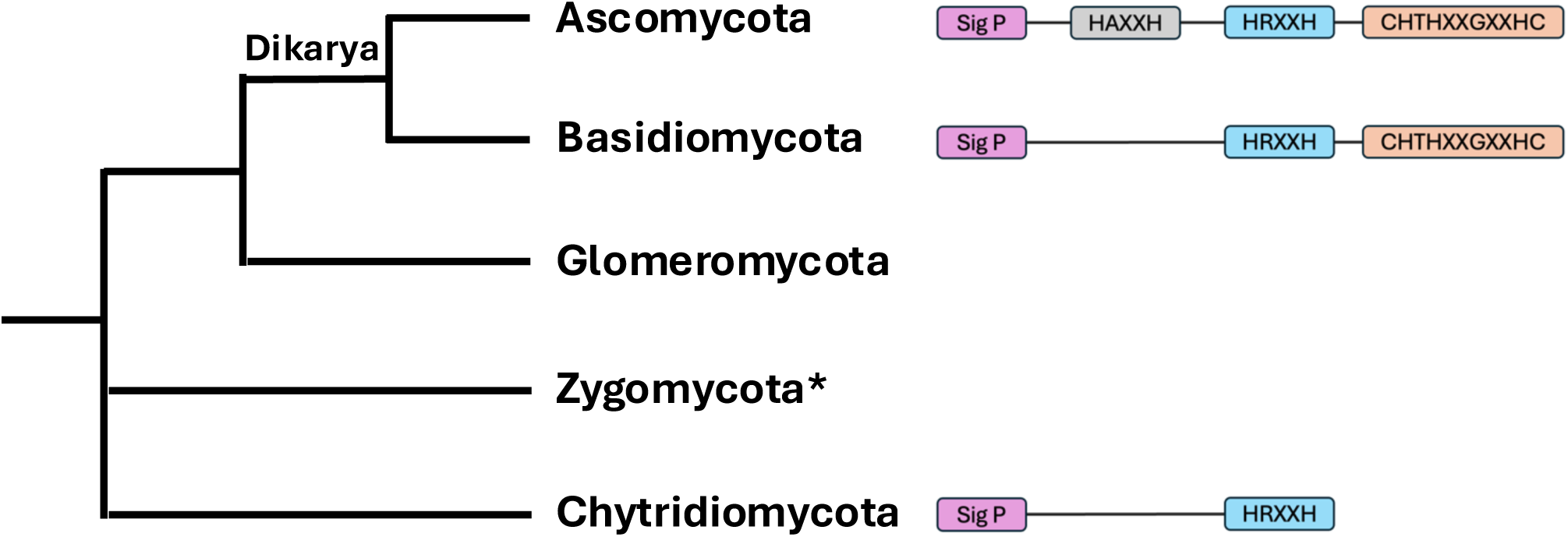
Simplified overview of fungal phylogeny and most common Pra1 signal peptide and predicted zinc binding motifs in each phyla. The asterisk denotes that Zygomycota is now two phyla, the Mucoromycota and Zoopagomycota.

This raises the intriguing question of what the zinc binding site in Pra1 looks like and how it compares to that of the catalytically active metalloproteases. From our cryo-EM structure of Pra1 prepared in the presence of 1 mM ZnSO_4_, we observe that the cryo-EM map is continuous amongst a cluster of three histidine residues (His178, His182 and His193), which all point towards each other, consistent with the coordination of a metal (Figure 2b). In this subunit (chain B), the centre of the zinc is equidistant to the centre of the τ nitrogen in the imidazole ring of each histidine residue. This distance is 2.3 Å. There is some extension of cryo-EM density in the metal binding site, likely indicating the presence of a water molecule to complete the tetrahedral coordination of the zinc. We did not apply symmetry when determining the cryo-EM map and thus we are able to observe that the precise distances from centre of the zinc to the centre of the τ nitrogen in the histidine residue vary by subunit (Extended Figure 4). For instance, in chain A, the distance from the centre of the zinc to the centre of the τ nitrogen of His 193 and His182 is 2.3 Å. In contrast, the distance from the centre of the zinc to the centre of the τ nitrogen of His 178 is 2.7 Å. These variations may be indicative of differential binding modes and might be important in the mechanism of zinc binding by the Pra1 hexamer.

**Figure 4.**
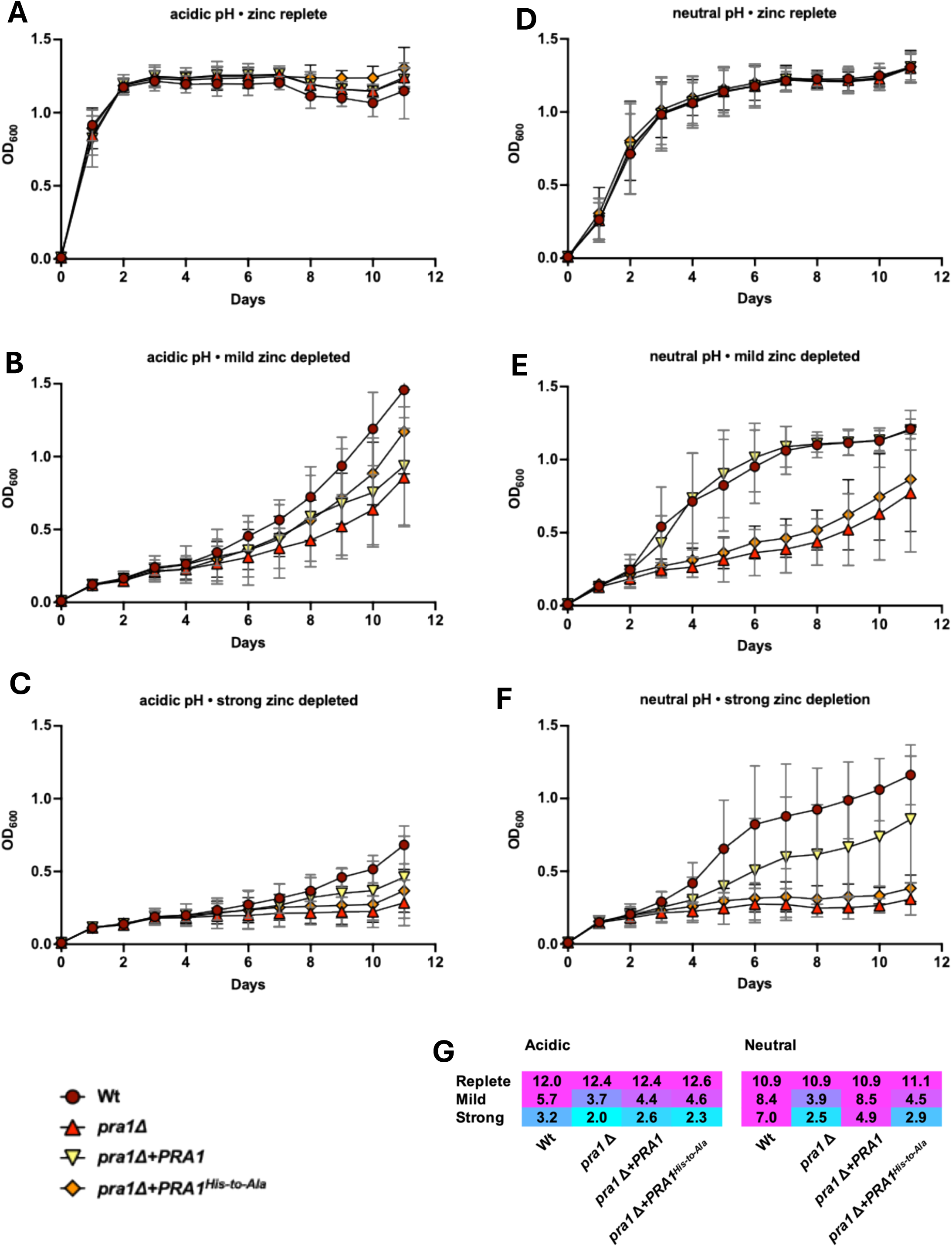
**The HRFWH motif is essential for *Candida albicans* growth under zinc restriction at neutral/alkaline pH**. **a-f**) Wild type (Wt), *PRA1* deletion mutant (*pra1*!1), *pra1*!1 expressing wild type *PRA1* (*pra1*!1+*PRA1*) and *pra1*!1 expressing HRFWH His-to-Ala variant (*pra1*!1+*PRA1^His-to-Ala^*) were cultured in acidic (unbuffered) or neutral/alkaline (buffered to pH 7.4) zinc free minimal medium (YNB-zinc-dropout; 0.5% glucose) supplemented with 1 μM zinc. To elicit mild- and strong- zinc depletion, media were supplemented with 0.5 mM or 2 mM EDTA, respectively. Cells were cultured at 30°C and 180 rpm. Growth was assessed by measuring OD_600_ for 11 days (n=5). **g**) The area under the curve was calculated for each growth curve and plotted with magenta colouring high and cyan colouring low measurements.

We observe the location of metal binding in Pra1 is conserved with that of the well characterised acid metalloprotease deuterolysin, by overlaying the Pra1 protomer with that of *Aspergillus oryzae* deuterolysin (Extended Figure 3b). The positions of Pra1-His182 and Pra1-His178 are conserved with those of deuterolysin-His132 and deuterolysin-His128, respectively. However, the catalytically active deuterolysin-Asp143 is substituted by Arg179 in Pra1, with the sidechain pointing away from the zinc binding site (Extended Figure 3b). In contrast to deuterolysin, this metal coordination site in Pra1 thus binds to zinc but is unlikely to be involved in proteolysis. In the context of overall Pra1 architecture, these six zinc binding sites are located on the outer face of the hexameric ring (Figure 2c).

To further compare the zinc-bound structure to Pra1 in the apo-state, we collected cryo-EM data on *C. albicans* Pra1 in MES pH 6.0, thereby ensuring protonation of histidine residues and release of any zinc due to electrostatic repulsion (Extended Figure 5). A comparison between the maps of Pra1 + zinc and Pra1 in MES pH 6.0 at the same contour level suggests that the His178, His182 and His193 triad no longer coordinate the metal ion under low pH conditions in the absence of zinc (Figure 2d). The lack of metal coordination, and the change in pH, does not lead to global conformational changes in the resolved portions of Pra1 (indeed a structural superposition of the hexamers in the presence and absence of zinc has an RMSD=0.175).

**Figure 5.**
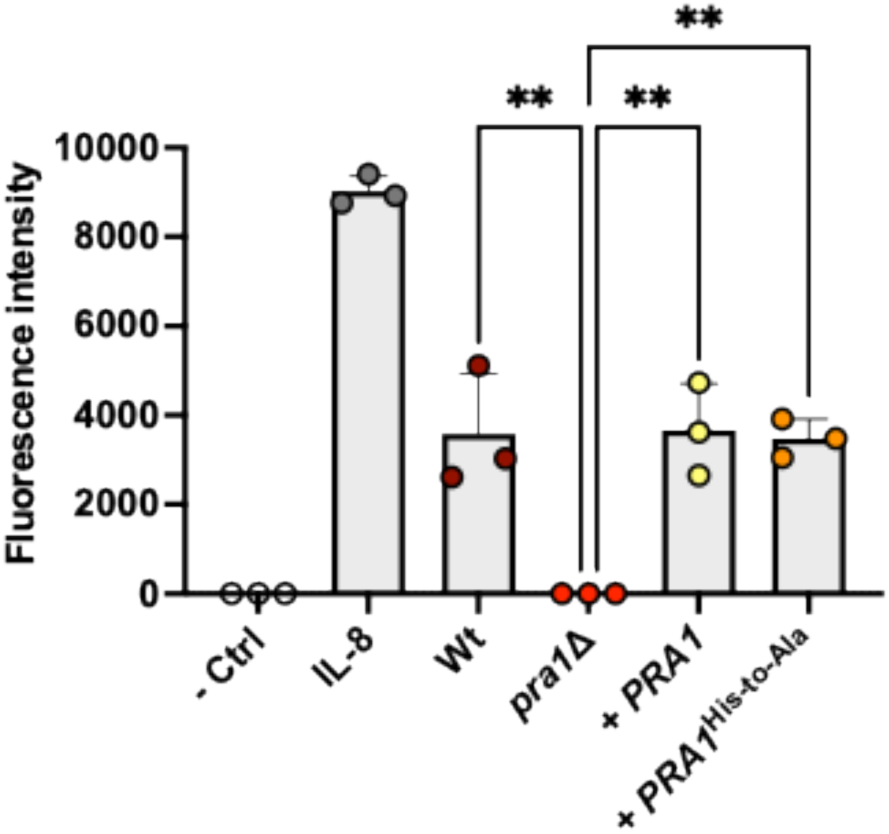
**Zinc binding motifs of Pra1 are not required for neutrophil recognition**. Wild type (Wt), *PRA1* deletion mutant (*pra1*Δ), *pra1*Δ expressing wild type *PRA1* (*pra1*Δ+*PRA1*) and *pra1*Δ expressing HRFWH/HARDH His-to-Ala variant (*pra1*Δ+*PRA1^His-to-Ala^*) were incubated in RPMI for five days. Culture filtrate, unconditioned media (RPMI) or media containing the neutrophil chemotactic factor, IL-8 (100 ng/ml), were added to the lower compartment of the chemotaxis plate. Freshly isolated human neutrophils were fluorescently labelled with Calcein and added to the upper compartment. Following 2 h incubation, neutrophil chemotaxis was determined by measuring the fluorescence intensity (at 485/530 nm) in the lower compartment (n=3). ** *P <* 0.01. ANOVA. Dunnett’s multiple comparison test amongst *C. albicans* culture filtrate samples.

This is the first structure of a zincophore and reveals an important role for the histidine residues at positions 178, 182 and 193 in zinc coordination. We next investigated the evolutionary and functional implications of these findings. The *PRA1* gene is ancient, having originated in an early fungal lineage and is present in most extant species^3,10^. As well as the HRXXH motif + His193 (residues 178-182, + 193) characterised in this study, *C. albicans* Pra1 contains an HAXXH motif (residues 68-72) and a C-terminal CHTHXXGXXHC motif (residues 289-299). We did not observe structural changes nor zinc binding at pH 6.0 nor pH 8.0 within proximity to residues 68-72. Whilst a short peptide spanning residues 289-299 does bind zinc, it is located at the end of what is predicted by Alphafold to be a long, unstructured C-terminus of the Pra1 protomer (Extended Figure 6) and is not resolved in our cryo-EM structures^7^.

**Figure 6.**
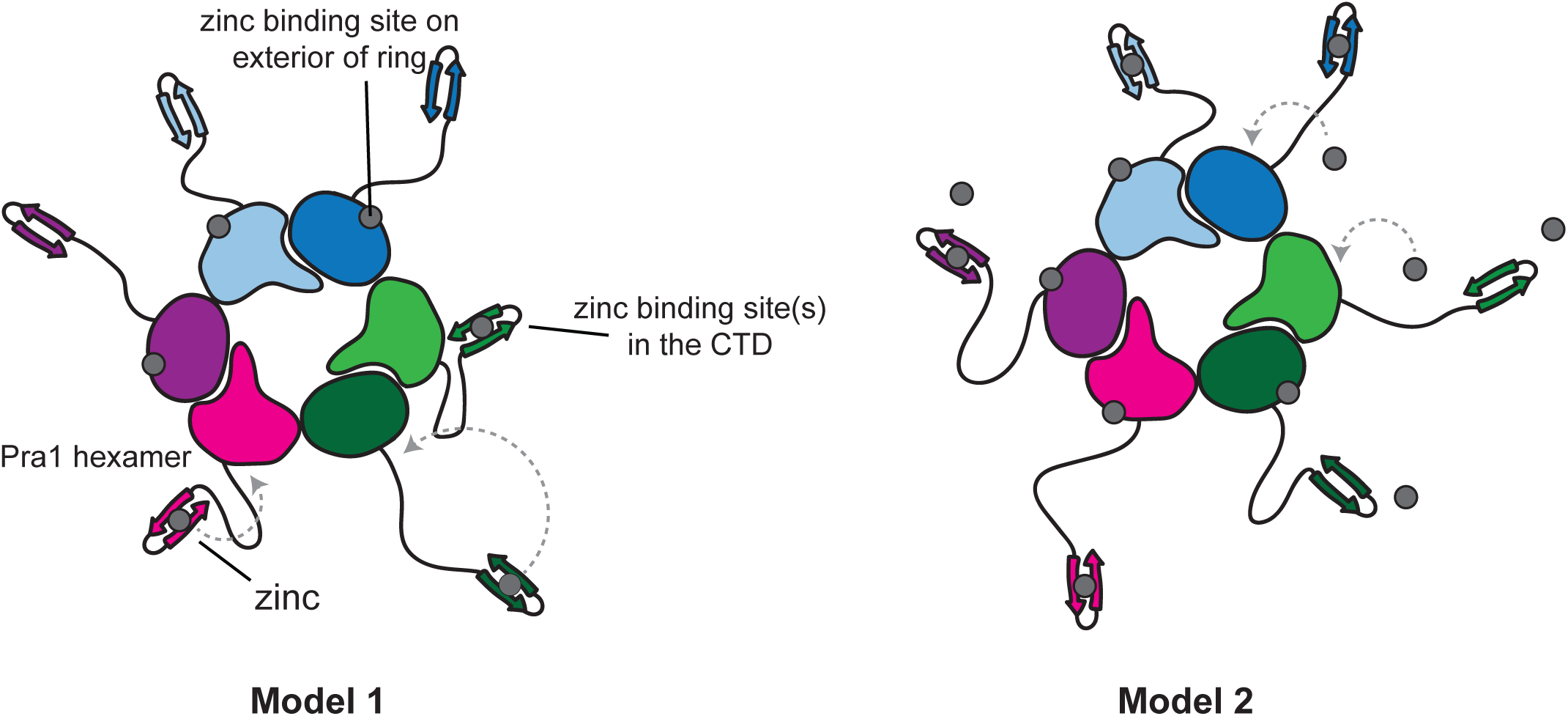
Possible models of molecular mechanisms of zinc scavenging by Pra1. Left, model 1 posits that the histidine triad functions as a zinc storage unit following capture of zinc by the C-terminal arms; right, direct capture of zinc by the histidine triads is illustrated in model 2.

BLASTp analysis of the *C. albicans* Pra1 primary amino acid sequence at NCBI identified hits only in fungi and in a single archaeon. Figure 3 shows a simplified overview of the phylogenetic relationship amongst fungal phyla. We observed 1336 hits in ascomycete fungi, 150 in basidiomycetes, two in chytridiomyetes and two in a Bathyarchaeota archaeon. Reciprocal BLASTp of the archaeal sequences identified orthologues in ascomycete fungi indicating possible horizontal transfer of the gene from fungi to this recently described archaeon.

Phylogenetic analysis of fungal Pra1 orthologues revealed the following. Most ascomycete fungi have orthologues with similar histidine-rich motifs as *C. albicans* Pra1. Several ascomycete fungi, including the medically relevant *Blastomyces percusus* have lost the C-terminal CHTHXXGXXHC motif (Extended Figure 7). A similar pattern was observed for the basidiomycetes, except none of the analysed sequences from this phylum possessed the HAXXH motif. Finally, the two chytrid proteins each possess the core HRXXH motif and lack both HAXXH and CHTHXXGXXHC. As we have reported previously, multiple fungal species have lost the Pra1 encoding gene^3^.

Because chytrids are an early diverging fungal phylum, it would appear that the origin of the Pra1 zincophore gene actually lies in a very early fungal lineage, probably upon mutation of the catalytic glutamic acid residue of a metalloprotease to arginine (Extended Figure 4). Following the divergence of the Dikarya from basal fungal lineages, the contemporary Pra1 gained its CHTHXXGXXHC motif – incidentally, the encoding gene formed a syntenic relationship with its receptor (Zrt101) in the Dikarya^3^. Finally, following divergence of the ascomycetes and basidiomycetes, the ascomycetes gained an additional HAXXH motif, and a small number of species lost the C-terminal CHTHXXGXXHC.

From this analysis, together with our structural observations (Figure 2b), we conclude that the 178-HRXXH-182 motif together with His193 represent the core zincophore domain of Pra1. We therefore created a modified *C. albicans* strain that expresses Pra1, but with the histidines of the HRXXH motif mutated to alanine (His178Ala and His182Ala) and compared this strain to *C. albicans* wild type (Wt), *PRA1* gene deletion mutant (*pra1*Δ) and genetic revertant (*pra1*Δ+*PRA1*).

When cultured in acidic minimal media without zinc-restriction, all *C. albicans* strains grew equally well (Figure 4a). Mild zinc depletion, elicited with 0.5 mM EDTA delayed the growth of all strains whilst strong zinc depletion (2 mM EDTA) effectively prevented growth (Figure 4b-c). In neutral/alkaline media without zinc restriction, all strains again grew equally well, albeit slightly slower than under acidic conditions (Figure 4d). This is because yeast-fungi generally grow better at acidic pH. Strikingly, upon mild zinc-restriction, wild type *PRA1*-expressing strains grew, whilst the *pra1*Δ null, and the Pra1 Histidine-to-Alanine variant both exhibited severely delayed growth and upon strong zinc depletion, growth of these strains was abolished (Figure 4e-f). From these growth assays together with their respective calculated areas (Area Under Curve, Figure 4g), it would appear that *C. albicans* specifically utilises the Pra1 HRXXH zincophore motif for growth under zinc limitation at neutral-alkaline pH.

As well as capturing zinc, we have recently shown that Pra1 is a major driver of the inflammatory immunopathology of vulvovaginal candidiasis^5^. However, it is not yet known whether these two functions are related. Although we did not detect zincophore activity of the additional HARDH motif in our structural analysis, this site has been implicated in zinc-binding and we reasoned that in a more complex host-pathogen interaction setting, it could also influence immune recognition^7^. Therefore, for these experiments, we created a second modified Pra1 in which the histidines of both HRXXH and HARDH motifs were substituted for alanine.

To test whether zincophore function is important for neutrophil recognition, we incubated our different *C. albicans* strains in RPMI tissue culture media for five days. Under this culture condition Pra1 is dispensable for growth and all strains grow to similar levels^5^. The fungal culture filtrates were added to the lower compartments of a chemotaxis assay plate; freshly isolated, labelled human neutrophils were then added to the upper compartment and allowed to migrate for 2 h. As we have observed previously, *C. albicans* wild type supernatant drove robust neutrophil recruitment, and this was blocked by deletion of *PRA1*. Genetic complementation of the *pra1*!1 mutant with a wild type copy of *PRA1* restored neutrophil recognition. Interestingly, genetic complementation of *pra1*Δ with the modified Pra1 variant fully restored neutrophil recognition to wild type levels (Figure 5). Therefore, the HRXXH zincophore motif is essential for *C. albicans* growth under zinc limitation, but not involved in neutrophil recognition.

## Discussion

Infections caused by *C. albicans* trigger a nutritional immune response by the host to inhibit pathogen growth. This results in environments where micronutrients such as zinc are scarce. To sequester zinc, *Candida albicans* deploys its zinc-scavenging machinery, a key component of which is the zincophore protein Pra1. In this study, we determined the cryo-EM structure of the *C. albicans* Pra1 both in the presence and absence of zinc, revealing a hexameric ring-like assembly. The 178-HRXXH-182 motif was previously described as a predicted zinc binding site^3^. Our study revealed a conserved histidine triad comprising His178, His182, His193. Our Pra1 structure determined in the presence of zinc showed cryo-EM density within this triad consistent with zinc coordination. Each Pra1 protomer hosts one triad, leading to six triads on the outside rim of the hexameric Pra1 assembly. We postulated that these triads are important for zinc sequestration and hence needed for *C. albicans* growth in zinc limited conditions. We made a variant *Pra1-His178Ala/His182Ala* strain and subsequently showed that this strain indeed has severe growth defects under low zinc conditions, specifically in a neutral-alkaline pH environment.

Our phylogenetic analysis of the Pra1 gene revealed its origin in a very early fungal lineage. The zincophore activity of Pra1 orthologues is likely also conserved. For instance, the *Aspergillus fumigatus* orthologue (AspF2) is required for growth under zinc limitation and has recently been shown to have zinc-binding properties^8,11^. Moreover, amino acid sequence alignment between *C. albicans* Pra1 and *A. fumigatus* AspF2 reveals that the histidine triad which we identified in Pra1 is conserved in AspF2.

An intriguing question arising from our study is that of the specific mechanism by which zinc binding by the histidine triad contributes to zinc scavenging by Pra1. The C-terminal residues, which harbour conserved histidine-rich regions, are not resolved in our Pra1 cryo-EM maps. This is most likely due to their inherent flexibility. Ser251, which is the most C-terminal residue that can be resolved in our cryo-EM maps, is pointing somewhat outward from the hexameric Pra1 assembly. It may be that the C-terminal tails radiate outward from the core hexameric ring (Figure 5). The relationship, if any, between these C-terminal zinc binding sites and the histidine triad remains to be defined. A possibility is that the C-terminal regions function like the arms of an octopus, foraging for zinc which is then transferred to the triads on the outside rim of Pra1, which act as storage units prior to uptake by *C. albicans* as hypothesised in Model 1 (Figure 6). Alternatively, the histidine triads may participate in zinc sequestration directly, as illustrated in Model 2 (Figure 6). It remains to be determined how the zinc-loaded Pra1 docks to the zinc transporter Zrt101 and facilitates zinc transfer. The work presented here is thus foundational for unravelling the structural basis of the Pra1-Zrt101 interaction in future work.

It has recently been shown that the Pra1 protein triggers the host inflammatory response in mucosal infections^5^. Our current study shows that the histidine triad is not involved in neutrophil recognition and thus is unlikely to contribute to the inflammatory response by the host. Our Pra1 cryo-EM data thus provide structural context for future work to investigate Pra1 interactions with the host proteome and subsequent therapeutics development.

## Materials and Methods

### Cloning, expression and purification of Pra1

The codon-optimised full-length *Pra1* (excluding the signal peptide) was cloned into the pCDNA3.0 plasmid with an N-terminal His-tag and a TEV protease cleavage site, using standard cloning techniques. Expi293 cells (50 mL culture) were transfected with the construct according to the manufacturer’s instructions, and the supernatant was collected five days post-transfection. Protein purification was performed using an ÄKTAgo system. The supernatant was loaded onto a 1 mL HisTrap excel column pre-equilibrated with 10 column volumes (CV) of 50 mM Tris (pH 8.0), 500 mM NaCl. After washing with 10 CV of the same buffer, the protein was eluted with 5 CV of 50 mM Tris (pH 8.0), 500 mM NaCl, and 400 mM imidazole. Eluted fractions were pooled and buffer-exchanged into TEV cleavage buffer (50 mM Tris, pH 8.0, 0.5 mM EDTA, 1 mM DTT). TEV cleavage was performed overnight using a 1:1000 molar ratio of TEV protease to target protein. The following day, the digestion mixture was passed over 0.5 mL Ni-NTA agarose resin (Thermo Scientific) using gravity flow to remove His-tagged TEV and other contaminants. The flow-through containing the cleaved protein was concentrated and subjected to size-exclusion chromatography (SEC) on a Superose 6 column. Peak fractions were collected, analyzed via SDS-PAGE, and pooled for further concentration and preparation for cryo-EM.

### Cryo-EM sample preparation, image collection and single-particle analysis

Pra1 was extensively buffered exchanged into either 20 mM MES pH 6.0, 150 mM NaCl or 20 mM Tris-HCL pH 8.0, 150 mM NaCl using spin filter concentrators with a 50 kDa cut-off (Amicon). Prior to sample freezing, BSA (Sigma) was added as an additive to 0.05%. For the Pra1:zinc sample, ZnSO4 was added to the sample to a final concentration of 1mM prior to freezing. Pra1 (3-4 ul) was applied to glow-discharged 1.2/1.3 Quantifoil 300 mesh carbon coated copper grids or lacey carbon copper grids (Electron Microscopy Services). The grids were blotted for 3.2 sec at 85% humidity and 20°C before plunge freezing into liquid ethane. Datasets were collected using a Titan Krios operated at an acceleration voltage of 300kV and the GATAN K3 direct electron detector coupled with the GIF quantum energy filter controlled using EPU software. Movies were recorded with a pixel size of 0.861 Å, and a dose rate of 2.1 e/Å^2^/frame. The program Warp was used to align movies, estimate the CTF and pick particles, using a 320 pixel box size ^12^. 2D classification, ab initio model non-uniform refinement were performed using cryoSPARC software^13^ . The data processing workflows for apo-state Pra1 and the Pra1:zinc complex are outlined in Extended Data Figures 1 and 5, respectively. Local resolutions of the cryo-EM maps were estimated using the programme ResMap ^14^. Model building was performed in UCSF Chimera 1.16 and Coot 0.9.8.7 ^15,16^. Models were depicted in figures using Pymol 2.5.4 and UCSF ChimeraX 1.7.1 ^17,18^. The AlphaFold model of *Candida albicans* Pra1 available on the AlphaFold Protein Structure Database was used as a starting model for manual building ^19,20^. The models were refined against the cryo-EM maps in Coot 0.9.8.7 and PHENIX ^16,21^. Data collection and refinement statistics are outlined in Table 1. Superpositions of atomic models and subsequent RMSD calculations were performed in Pymol 2.5.4 using the align algorithm ^17^.

**Table 1.** Cryo-EM data collection and refinement statistics.

|  |  |  |
| --- | --- | --- |
|  | Pra1:zinc complex<br>EMD-xxxxx<br>PDB-xxxx | Pra1<br>EMD-xxxxx<br>PDB-xxxx |
| **Data collection and processing** |  |  |
| Microscope | Titan Krios | Titan Krios |
| Camera | K3/counting | K3/counting |
| Magnification | 105,000 | 105,000 |
| Energy filter | Gatan | Gatan |
| Energy filter slit width (eV) | 20 | 20 |
| Collection software | EPU | EPU |
| Voltage (kV) | 300 | 300 |
| Cumulative exposure (e-/Å<sup>2</sup>) | 63.2 | 63.2 |
| Exposure rate (e-/Å<sup>2</sup>/frame) | 2.1 | 2.1 |
| Defocus range (μm) | -1.0 – -2.4 | -1.0 – -2.4 |
| Pixel size (Å) | 0.861 | 0.861 |
| Symmetry imposed | C1 (none) | C1 (none) |
| Number of micrographs | 10,785 | 6,886 |
| Initial particle images (no.) | 3,486,720 | 1,025,587 |
| Final particle images (no.) | 1,102,286 | 607,405 |
| 0.143 FSC half map masked (Å) | 2.53 | 2.85 |
| 0.143 FSC half map unmasked(Å) | 3 | 3.3 |
| **Refinement** |  |  |
| Refinement package | Phenix | Phenix |
| Initial model used (PDB code) | The AlphaFold model of *C. albicans* Pra1 as a starting model for manual building | Pra1:zinc model, followed by manual building |
| Model resolution range (Å) | 2.25-3.25 | 2.5-3.5 |
| Model composition |  |  |
| Non-hydrogen atoms | 10626 | 10620 |
| Protein residues | 1326 | 1326 |
| Ligands | ZN:6 | n/a |
| CC map vs. model (%) | 0.90 | 0.90 |
| R.m.s. deviations |  |  |
| Bond lengths (Å) | 0.002 | 0.004 |
| Bond angles (°) | 0.508 | 0.543 |
| Validation |  |  |
| MolProbity score | 1.04 | 1.29 |
| Clash score | 0.54 | 1.22 |
| Poor rotamers (%) | 1.84 | 2.11 |
| Ramachandran plot |  |  |
| Favored (%) | 97.11 | 96.73 |
| Allowed (%) | 2.89 | 3.27 |
| Outliers (%) | 0 | 0 |
| C-beta deviations | 0 | 0 |
| EMRinger Score | 3.36 | 3.19 |
| CaBLAM outliers (%) | 2.23 | 2.61 |

**Table 2.**
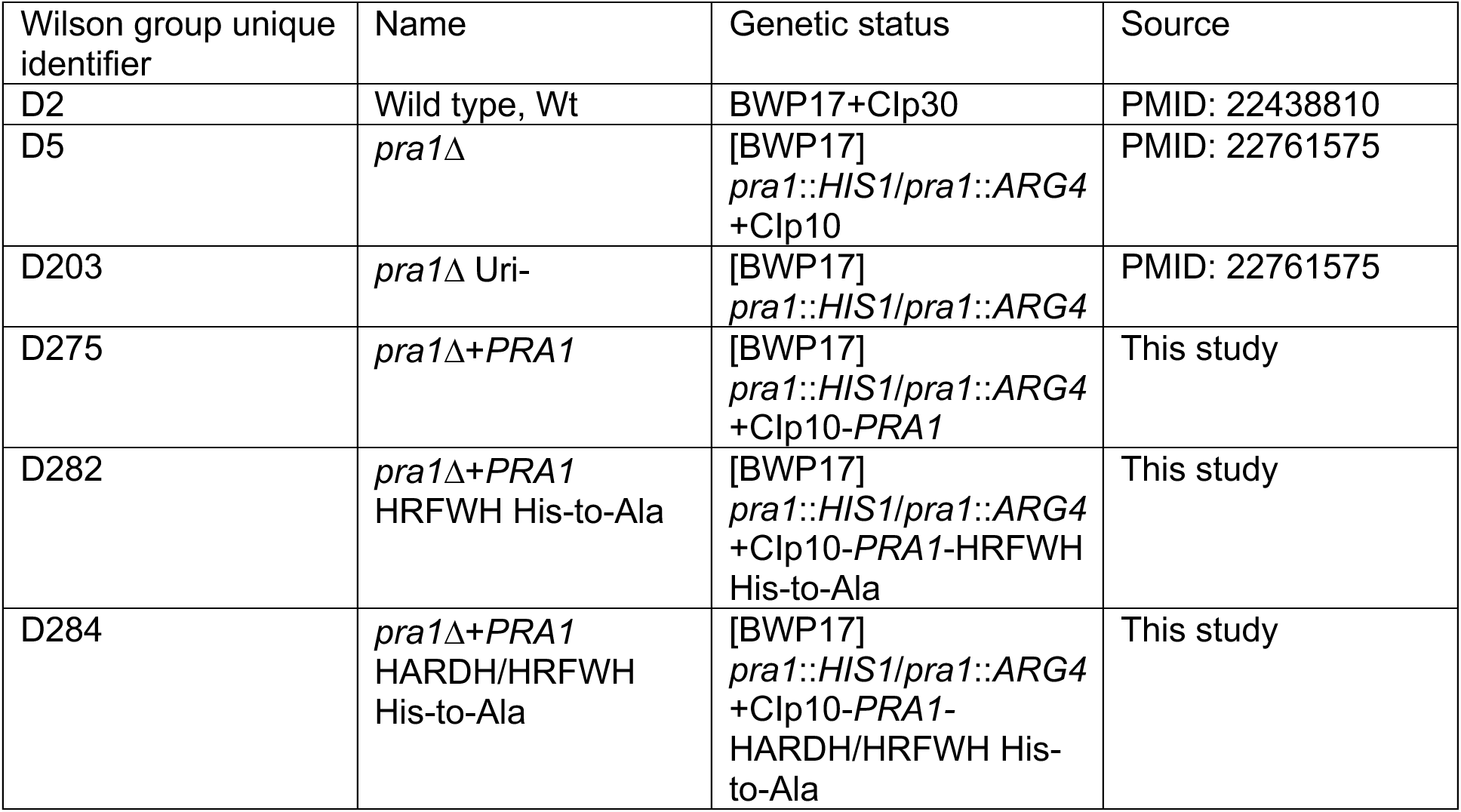
Strains of *Candida albicans* used in this study.

| Wilson group unique identifier | Name | Genetic status | Source |
| --- | --- | --- | --- |
| D2 | Wild type, Wt | BWP17+Clp30 | PMID: 22438810 |
| D5 | *pra1*Δ | [BWP17]<br>*pra1::HIS1/pra1::ARG4*<br>+Clp10 | PMID: 22761575 |
| D203 | *pra1*Δ Uri- | [BWP17]<br>*pra1::HIS1/pra1::ARG4* | PMID: 22761575 |
| D275 | *pra1*Δ+*PRA1* | [BWP17]<br>*pra1::HIS1/pra1::ARG4*<br>+Clp10-*PRA1* | This study |
| D282 | *pra1*Δ+*PRA1*<br>HRFWH His-to-Ala | [BWP17]<br>*pra1::HIS1/pra1::ARG4*<br>+Clp10-*PRA1*-HRFWH<br>His-to-Ala | This study |
| D284 | *pra1*Δ+*PRA1*<br>HARDH/HRFWH<br>His-to-Ala | [BWP17]<br>*pra1::HIS1/pra1::ARG4*<br>+Clp10-*PRA1*-<br>HARDH/HRFWH His-to-Ala | This study |

### Pra1 variant construction

*PRA1* alleles containing HRFWH and HARDH/HRFWH His-to-Ala coding variants were synthesised (GeneArt) consisting of the upstream intergenic region, open reading frame of allele B of *PRA1* and 100 base pairs downstream sequence (*Candida* Genome Database)^22^. However, *C. albicans* transformed with these constructs did not express Pra1 protein. Therefore, versions containing the whole allele (entire upstream and downstream intergenic regions) were generated. To do this, the wild type *PRA1* was amplified from *C. albicans* genomic DNA using primers (AN-oli56) SalI_pPRA1_F and (AN-oli57) PRA1_UTR554_MluI_R, cloned into TOPO vector, confirmed by Sanger sequencing and then subcloned into CIp10 for integration (Murad 2000 Yeast). We selected a CIp10-*PRA1* harbouring allele B of *PRA1* for subsequent modification. The resulting CIp10-*PRA1* plasmid was digested with *Sal*I and *Nde*I to excise the first 1,192 base pairs of insert, containing the upstream intergenic region and first 684 base pairs of *PRA1* coding sequence. The remaining backbone, 3’ of the gene and downstream intergenic region was purified by gel extraction. In parallel, The first 1,192 base pairs of HRFWH and HARDH/HRFWH His-to-Ala coding variants were excised from the synthetic constructs, purified by gel extraction, and independently cloned in the CIp10-PRA1 backbone fragment. Each (*PRA1* wild type, *PRA1*-HRFWH His-to-Ala and *PRA1*-HARDH/HRFWH His-to-Ala variants) of the plasmids were linearised and integrated at the *RPS1* locus of the *pra1*Δ uridine auxotroph as previously described (PMID: 22761575).

### Growth assays

*C. albicans* strains were maintained on YPD agar plates (1% yeast extract; 2% peptone; 2% glucose; 2% agar). A colony was inoculated into SD minimal medium (1X yeast nitrogen base; 2% glucose) and incubated at 30°C and 180 rpm. Precultures were then washed twice in milliQ water and used to inoculate culture media in 96 well plates to OD600 = 0.01. In all cases, the basal media was zinc-free minimal media supplemented with 1 μM zinc (1X YNB zinc- dropout (Formedium); 0.5% glucose; 1 μM ZnSO_4_). To buffer to neutral/alkaline pH, 80 mM HEPES pH 7.4 was added. To elicit zinc restriction, EDTA was added to a concentration of 0.5 or 2 mM. The complete media, including EDTA was always first prepared before the addition of *C. albicans* cells. Plates were incubated at 30°C and 180 rpm and OD_600_ measured daily.

### Neutrophil chemotaxis assay

*C. albicans* strains were cultured for 24 h in SD minimal media at 30°C, 200 rpm, washed twice in milliQwater, and resuspended in RPMI without phenol red at 1 x10^6^ cells/ml. These were then distributed in triplicate in a 24-well plate, 1 mL per well, and incubated at 24°C for 5 days, 130 rpm shaking. Conditioned supernatant was collected, filter sterilized (0.2 µm filter, Fisher) and added to the basal compartment of the chemotaxis chamber.

Human peripheral blood neutrophils (PMN), obtained from healthy volunteers, were separated by density gradient centrifugation on FicollPaque^TM^ Plus, followed by the hypotonic lysis of erythrocytes using a 1X red blood cells lysis buffer (10X recipe: NH_4_Cl 1.55 M, NaHCO_3_ 120 mM, EDTA 1mM, in sterile ddH_2_O) Using this protocol, we demonstrated the isolation of a neutrophil population with more than 98% purity using Flow Cytometry (PMID: 38055800). The purity of human neutrophils, before each chemotaxis test, was always checked using a Wright-Giemsa Stain (abcam) following the kit instruction. The slides were prepared using a Cytospin 4 centriguge (Thermo Scientific) and then evaluated microscopically using EVOS M5000 at 40x magnification.

Neutrophils were then washed twice with PBS and resuspended in 5 ml RPMI-1640 without phenol red containing 10% heat-inactivated FBS. The cells were then incubated with Calcein AM, fluorescent cell permeable derivative of calcein (Sigma Aldrich) (5 µg / ml) for 30 min at 37°C + 5 % CO_2_. After the incubation, the neutrophils were washed twice with PBS, counted, and resuspended in the same complete medium at 5 x 10^6^ / ml.

Human neutrophil chemotaxis was measured using a 96-well chemotaxis chamber with 3.2 mm diameter filter with membrane porosity of 5 µm (Neuro Probe). Labelled human neutrophils (5 x 10^6^/ml) were transferred into ChemoTx filters placed in the 96-well plates containing 300 μl of either medium alone (RPMI without phenol red, negative control), media containing recombinant IL-8 (100 ng/ml) (Biochem, positive control) or an appropriate dilution of *Candida* culture filtrate supernatant in the same media. The chamber was incubated for 2 h at 37°C + 5% CO_2_. Following incubation, the non-migrating cells on the top of the filter were removed by gently wiping the filter and the cells that had migrated into the bottom chamber were measured by fluorescence signal (excitation, 485 nm; emission, 530 nm) using a Tecan Spark plate reader.

## Acknowledgements

JLS would like to thank Prof. Hiro Furukawa for advice, lab resources and encouragement, as well as Dr. Dennis Thomas and Dr. Ming Wang for managing the cryo-EM and computing facilities, respectively, at Cold Spring Harbor Laboratory. JLS is supported by the Research Council of Finland. DW supported by the MRC Centre for Medical Mycology at the University of Exeter (MR/N006364/2 and MR/V033417/1), the NIHR Exeter Biomedical Research Centre (NIHR203320), and the Wellcome Trust (214317/A/18/Z).

Additional work may have been undertaken by the University of Exeter Biological Services Unit. The views expressed are those of the author(s) and not necessarily those of the NIHR or the Department of Health and Social Care.

Molecular graphics for several figures was performed with UCSF ChimeraX, developed by the Resource for Biocomputing, Visualization, and Informatics at the University of California, San Francisco, with support from National Institutes of Health R01-GM129325 and the Office of Cyber Infrastructure and Computational Biology, National Institute of Allergy and Infectious Diseases.

## Data Availability

The final cryo-EM maps have been deposited in the Electron Microscopy Data Bank under accession codes: EMD-xxxxx and EMD-xxxxx. The final models have been deposited in the Protein Data Bank under the accession codes: PDB-xxxx and PDB-xxxx. The PDB entry 1EB6 was used for strcutural superposition.

