## Supplemental_Material for "Structural insights into mechanisms of zinc scavenging by the *Candida albicans* zincophore Pra1"

#### Extended Data Figures and Figure Legends

**Extended Figure 1. Workflow of cryo-EM structure determination of *C.albicans* Pra1 in the presence of zinc.** a) A representative micrograph. b) Representative 2D classes. c) Overview of the Initial motion correction, CTF estimation and particle picking were performed in Warp, other steps were performed in cryoSPARC [1,2]. d) The Fourier Shell Correlation graph. e) A local resolution map calculated using ResMap [3].

**Extended Figure 2. The cryo-EM map of *C. albicans* Pra1 exhibits extensive N-linked glycolysation.** Residues N48, N89, N135 and N208 are glycosylated. Two glycans are in close proximity to the inter-dimer interface, whereas six glycans are in close proximity to the intra-dimer interface.

**Extended Figure 3. *C. albicans* Pra1 overlaid with the metalloprotease deuterolysin (pdb:1eb6).** a) The overall fold of a Pra1 subunit is similar to that of deuterolysin (PDB code: 1EB6). b) The canonical catalytically active aspartic or glutamic acid in HEXXH+D type metalloproteases is not conserved in *C. albicans* Pra1. Deuterolysin D143 is in the same position as Pra1 R179, but the R179 sidechain not pointing towards the zinc coordination site. For clarity, only the zinc from PBD-1EB6 is shown.

**Extended Figure 4. The positioning of the histidine residues in the metal binding site varies by subunit.** The distance from the centre of the zinc to the centre of the τ nitrogen of His 193 and His 182 is 2.3 Å. The distance from the centre of the zinc to the centre of the τ nitrogen of His 178 is 2.7 Å. The distances are indicated by the dashed lines.

**Extended Figure 5. Workflow of cryo-EM structure determination of *C.albicans* Pra1 in the absence of zinc.** a) A representative micrograph. b) Representative 2D classes. c) Overview of the Initial motion correction, CTF estimation and particle picking were performed in Warp, other steps were performed in cryoSPARC [1,2]. d) The Fourier Shell Correlation graph. e) A local resolution map calculated using ResMap [3].

**Extended Figure 6. AlphaFold predicted structure of a Pra1 monomer**. Using a confidence score from 1-100, the core of the protein is predicted with very high confidence (>90), the C-terminal motif with high confidence (>70) and the unstructured region with low (>50) to very low confidence (<50) [4,5].

**Extended Figure 7.** Sequence alignment of Deuterolysin from *Aspergillus oryzae* and Pra1 orthologues basidiomycetes (*Ustilago maydis*, *Cryptococcus depauperatus*), chytrids (*Spizellomyces punctatus*) and ascomycetes (*Blastomyces percusus*, *Candida albicans* and *Aspergillus fumigatus*). Calculated using ESPript - https://espript.ibcp.fr [6].

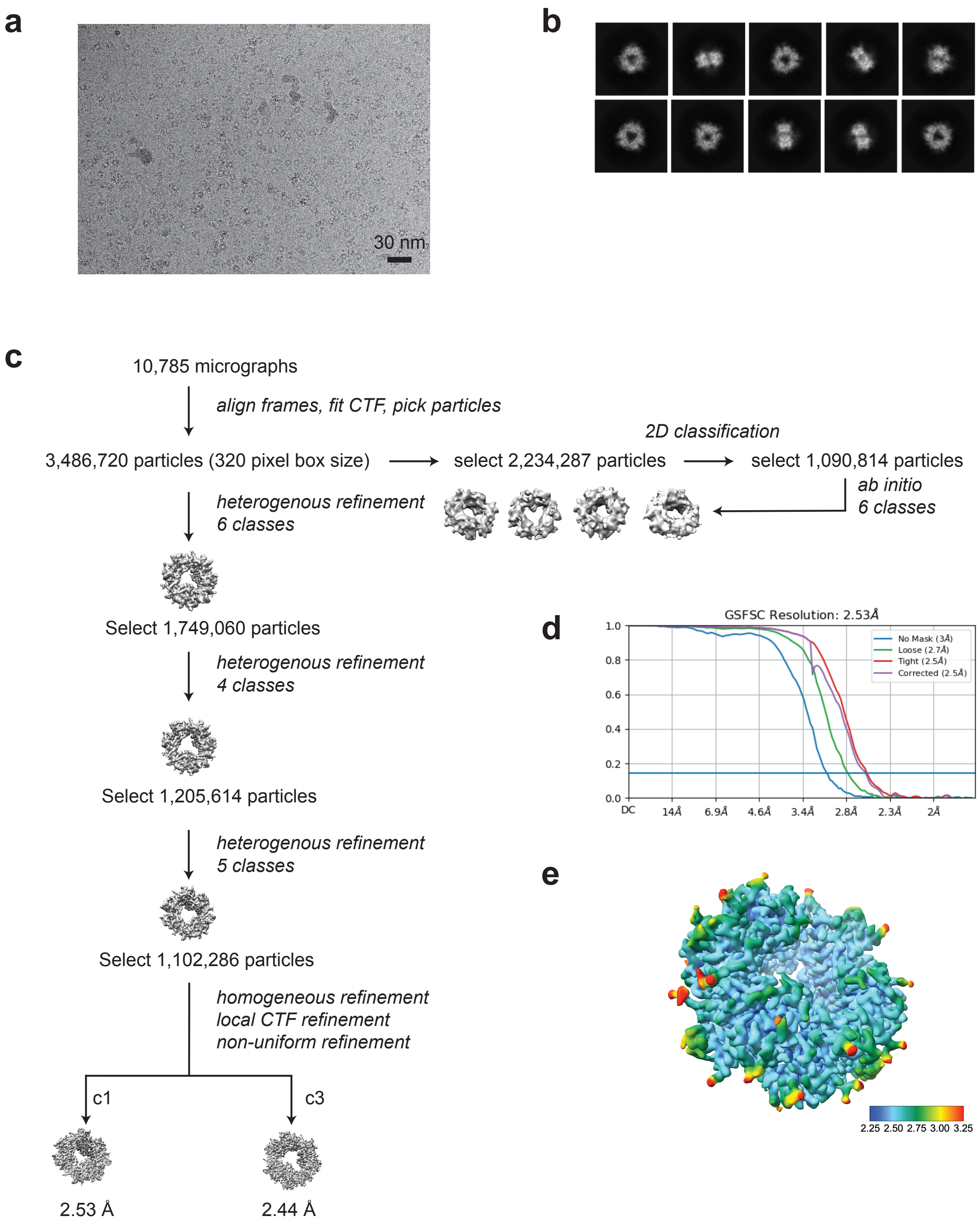


### Extended Figure 1

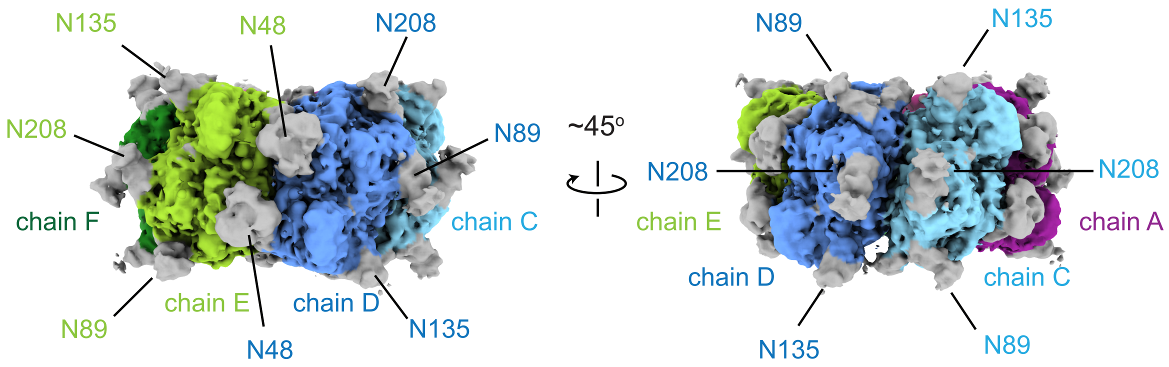


### Extended Figure 2

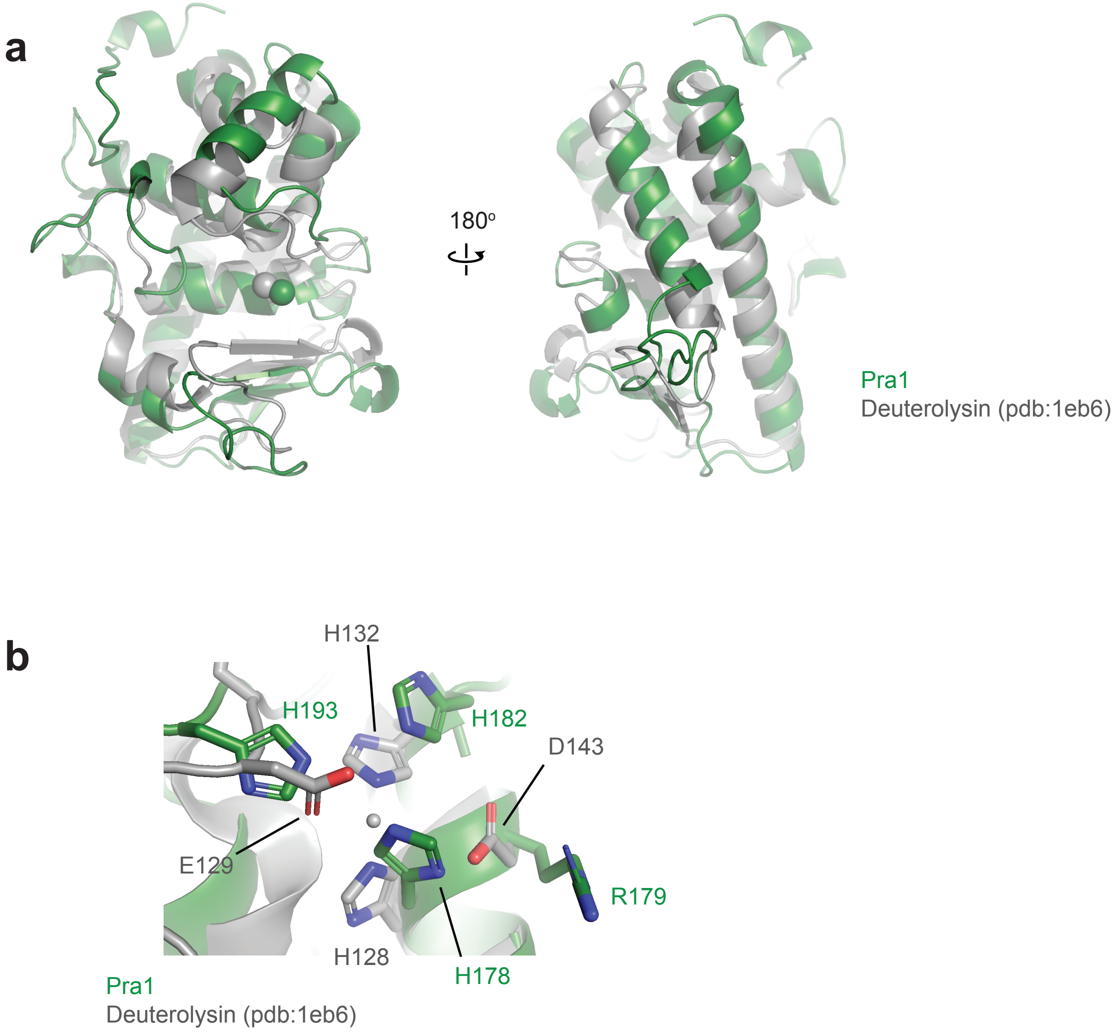


**Extended Figure 3**

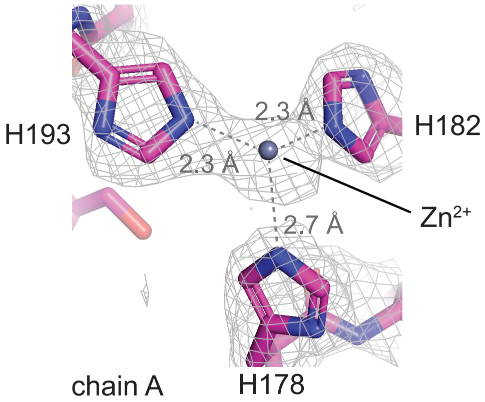


#### Extended Figure 4

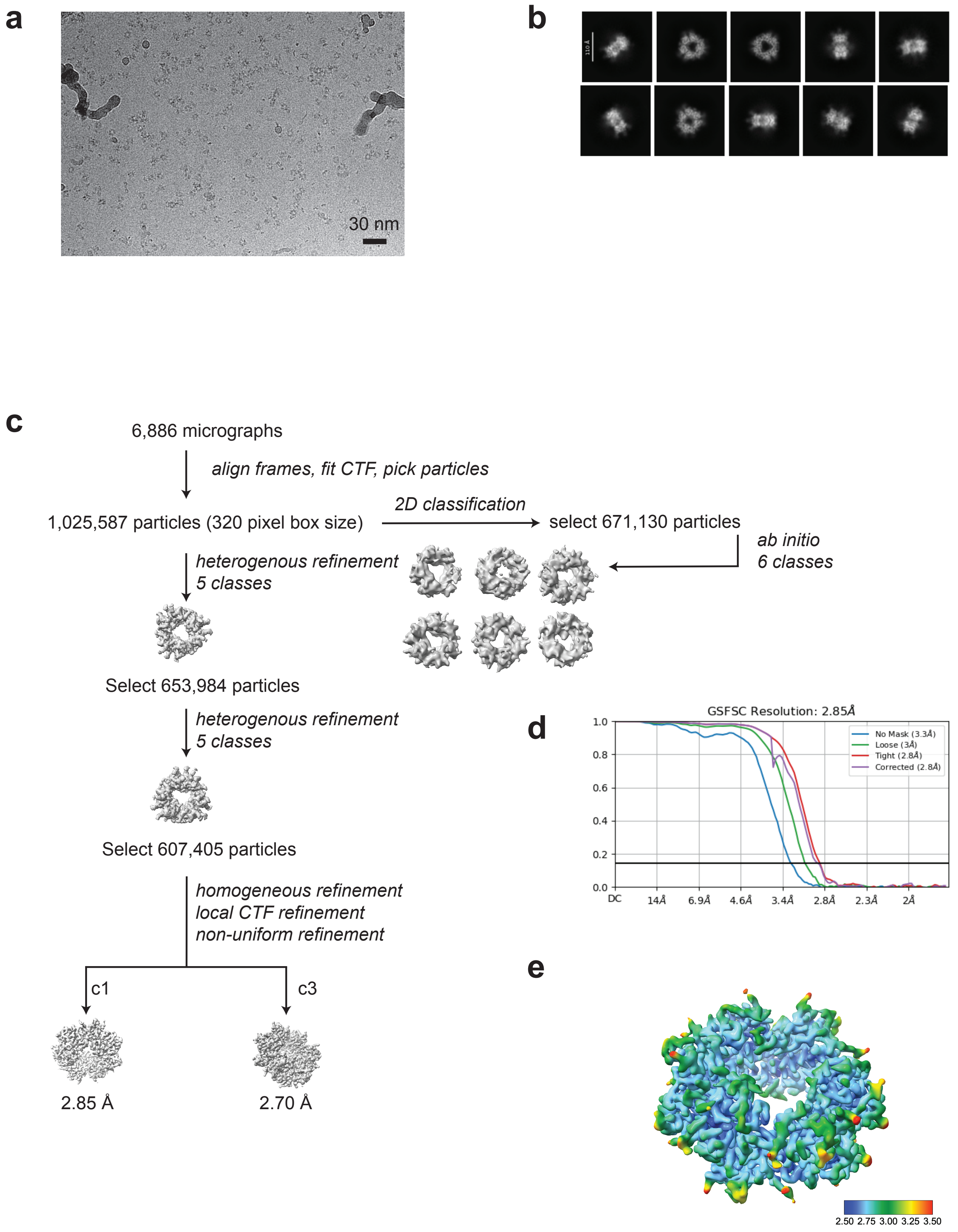


### Extended Figure 5

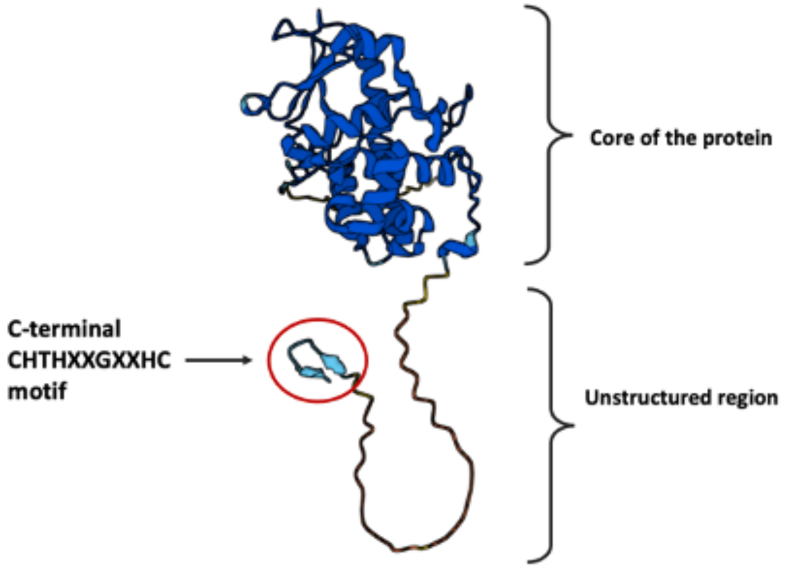


**Extended Figure 6**

```
Deuterolysin                    VTKALSQLTRRTEVTDCKGDAESSLTTALSNAAKLANQAAEAA--ESGDESKFEEYFKTT    222
Ustilago_maydis                 -----QIDSVWRIHESCNGTQRAQISSGIDDMKKLAHN-SINHILNYPKDEFFIKYFGQD    93
Cryptococcus_depauperatus       ------YTSDINIHESCNATQRRMLDKALSDAFEVASFAKEYARTNGPADPVFQKYFGKD    80
Spizellomyces_punctatus         -RRQLPPSTRVQMHSSCSASQKQTLNKALDDMNKLTIHATKRILDKTYEDPVYQTYFGNG    81
Rhizophlyctis_rosea             LSNGKGYNADVTIHASCKGTQERALTKALGEMNDLAKIAANRILKYGSEDELYKKYFGDG    91
Blastomyces_percursus           RKCGYGMMDPYPIHDSCNATERRMISRGLDDAITLAAHARDHVLKFGHDSSLYRKYFGNA    133
Candida_albicans_Pra1           ------WVKGFPIDSSCNATQYNQLSTGLQEAQLLAEHARDHTLRFGSKSPFFRKYFGNE    90
Aspergillus_fumigatus_AspF2     ------AVTSFPIHSSCNATQRRQIEAGLNEAVELARHAKAHILRWGNESEIYRKYFGNR    103
                                               .*..     :  .: :   ::             .  :  **

Deuterolysin                    DQQTRTTVAERLRAVAKEAGSTSGGSTTYHCNDPYGYCEP-NVLAYT---LPSKNEIANC    278
Ustilago_maydis                 ADPA-PVVGYYVEL-----VYGNKGDALLRCDNPDDNCRL-PEWNGHWRGNNATAETVIC    146
Cryptococcus_depauperatus       ADSYTQVIGIWDAF-----LTGNKEGVILRCDNPDGNCAQ-KGFNGHWRGNNATSETVIC    134
Spizellomyces_punctatus         ES-A-TVVGYYGIL-----TAGK-----VFFSDVDNKCSQ-PGWAGHWRGEVAPLETVIC    128
Rhizophlyctis_rosea             EA-A-TVLGYYKTI-----LYGNKPGVLFRCDNIDNKCHQ-EGWAGHWRTEIAPLETVIC    143
Blastomyces_percursus           PT-S-NVIGNLARI-----VDGNRKKTLFRCDDPDGNCKRIPTYGGHWRGENATDETVIC    186
Candida_albicans_Pra1           TASA-EVVGHFDNV-----VGADKSSILFLCDDLDDKCKN-DGWAGYWRGSNHSDQTIIC    143
Aspergillus_fumigatus_AspF2     PT-M-EAVGAYDVI-----VNGDKANVLFRCDNPDGNCAL-EGWGGHWRGANATSETVIC    155
                                      .:.             .        .:  . *                 :   *

Deuterolysin                    DIYYS--ELPPLAQKCHAQDQA----------TTTLHEFTHAPGVYQ-PGTEDLGYGYDA    325
Ustilago_maydis                 ELSYV--TRRPLEKLCSAGFQLGTDNPSLYFGADLMHRAFHVPEFVH-EKIHHYADSYAD    203
Cryptococcus_depauperatus       DLSYT--SRIYNEAFCMSGFQLASQKPSTYWSVDLIHRFFHVPAVTN-GLVGHFAEDYAS    191
Spizellomyces_punctatus         PLSFNTTARKPLSALCQDGDRFSQHKTNYFLSSDLMHRLFHVPLINPEERVDHYASNYTE    188
Rhizophlyctis_rosea             PLSFT-DARKPLSAICKNGNKISAVKSNYFFASDLMHRLFHVPLINPGERVDHFAGNWTE    202
Blastomyces_percursus           ELSYK--TRLYLEHFCMGGYTVAKSPRNTYFGLDMMHRLYHMPAIGE-NHVGHFADTYND    243
Candida_albicans_Pra1           DLSFV--TRRYLTQLCSSGYTVSKSKTNIFWAGDLLHRFWHLKSIGQ-LVIEHYADTYEE    200
Aspergillus_fumigatus_AspF2     DRSYT--TRRWLVSMCSQGYTVAGSETNTFWASDLMHRLYHVPAVGQ-GWVDHFADGYDE    212
                                   :           *                   :*.  *   .       . .  :

Deuterolysin                    ------------------------------------------------------------    352
Ustilago_maydis                 APTST--------PAA----------------------------------TPTPSAASSD    278
Cryptococcus_depauperatus       ATSSNA----GAASASVPPPPAESTGCTLHGDHYHCTGPATPTTQAAEVHDTPASGGDKD    307
Spizellomyces_punctatus         ------------------------------------------------------------    227
Rhizophlyctis_rosea             ------------------------------------------------------------    241
Blastomyces_percursus           ATQ-PT-------KG-TPTVP---------------------------HPPTPTADVPRV    321
Candida_albicans_Pra1           -SG-SDSGASSTASSSHQHTD---------------------------SNPSATTDANSH    288
Aspergillus_fumigatus_AspF2     ASTSTSSSSSGSGSGATTTPT---------------------------DSPSATIDVPPN    302

Deuterolysin                    -------------- 352
Ustilago_maydis                 CHTHADGSIHCGTH 292
Cryptococcus_depauperatus       CHTHADGTVHCV-- 319
Spizellomyces_punctatus         -------------- 227
Rhizophlyctis_rosea             -------------- 241
Blastomyces_percursus           G------------- 322
Candida_albicans_Pra1           CHTHADGEVHC--- 299
Aspergillus_fumigatus_AspF2     CHTHEGGQLHCT-- 314
```

### Extended Figure 7
